# Systematic analysis of naturally occurring missense mutations in human manganese transporters: prediction and structural insights

**DOI:** 10.1101/2025.09.09.675244

**Authors:** Ryan Hu, Jian Hu

## Abstract

Manganese (Mn) homeostasis in humans is tightly regulated by the transporters ZIP8, ZIP14, and ZnT10, and pathogenic mutations in these proteins cause systemic Mn dysregulation, leading to severe disorders in multiple systems. Here, we performed a systematic survey of naturally occurring missense variants in these transporters. Pathogenicity was assessed with two widely used computational tools, CADD and AlphaMissense (AM), and results were integrated with AlphaFold-predicted structural models. Although the prediction methods showed general agreement, substantial discrepancies were observed, with CADD tending to overpredict deleteriousness and AM failing to identify a portion of confirmed pathogenic variants. Structural mapping revealed that predictions were more accurate for amino acid substitutions in structured and buried residues than for solvent-exposed loops and flexible regions. Mechanistic insights further highlighted variants at functionally important regions as critical. Our study helps prioritize Mn transporter variants for future experimental validation and provides a framework for bridging computational predictions with structure-guided mechanistic studies.

## Introduction

Manganese (Mn) is an essential trace element that functions as a cofactor for enzymes involved in antioxidant defense, neurotransmitter synthesis, and energy metabolism, making tight systemic and cellular Mn homeostasis critical for human health.^1, 2^ In humans, three genetically and functionally distinct transporters, i.e. the Zrt-/Irt-like protein (ZIP) family members ZIP8 (SLC39A8) and ZIP14 (SLC39A14), and the zinc transporter (ZnT) family member ZnT10 (SLC30A10), are central determinants of Mn absorption, distribution, and excretion.^3^ Divalent metal transporter 1 (DMT1, or SLC11A2) also contributes to Mn uptake, especially under conditions of iron deficiency when its expression is induced.^4, 5^

ZIP8, ZIP14 and ZnT10 act cooperatively to maintain systemic Mn homeostasis.^3^ Functioning as a metal importer, ZIP8 supplies cytosolic Mn that is required for Mn-dependent enzymes (including Golgi glycosyltransferases). In the liver, ZIP8 at the apical canalicular membrane of hepatocytes facilitates the uptake of Mn from bile into hepatocytes, preventing excessive loss of Mn through biliary excretion. ZIP14 is highly expressed in hepatocytes and at interfaces with the circulation where it mediates uptake of Mn from blood into the liver (and other tissues) and ZIP14-dependent hepatic Mn uptake is a key step in whole-body Mn clearance. ZnT10 is a dedicated Mn efflux transporter localized to the hepatocyte and enterocyte surface (and other cell types) that exports excess Mn into bile and the intestinal lumen, representing the principal Mn excretion pathway.^6^ Because of their crucial functions in systemic Mn homeostasis, perturbation of any of these Mn transporters shifts tissue Mn fluxes and yields the distinct biochemical and clinical phenotypes observed in human diseases. Indeed, loss-of-function (LOF) variants in Mn transporters are known to cause various clinically syndromes. LOF mutations in *slc39a8* cause SLC39A8-congenital disorder of glycosylation (SLC39A8-CDG, or CDG type 2n), a congenital disorder of glycosylation associated with Mn deficiency.^7-11^ This multisystem infantile-onset syndrome is characterized by global developmental delay, intellectual disability, hypotonia, movement abnormalities (including dystonia), seizures and variable dysmorphic features. By contrast, mutations in *slc39a14* produce an autosomal recessive hypermanganesemia with early-onset parkinsonism-dystonia.^12^ Pathogenic variants of SLC30A10 (ZnT10) cause hypermanganesemia with dystonia, polycythaemia and chronic liver disease (sometimes termed SLC30A10-related manganism), a disorder featuring neurologic movement disorder (dystonia/parkinsonism), very high serum Mn, hepatic fibrosis/cirrhosis and erythrocytosis.^13-15^

Large-scale sequencing projects and curated databases such as UniProt and gnomAD have catalogued millions of naturally occurring human genetic variants.^16, 17^ Yet, the vast majority of these entries remain poorly annotated, with limited or no experimental evidence regarding their functional consequences. This lack of annotation is a pervasive problem across most genes, where only a small fraction of variants can be confidently classified as pathogenic or benign. Instead, most are designated as “variants of uncertain significance” (VUS), reflecting the gap between rapid advances in variant discovery and the slower pace of functional characterization. This gap hinders both clinical interpretation and mechanistic studies, especially in cases where even subtle amino acid substitutions may disrupt protein activity, interactions, or localization in ways that prediction algorithms cannot reliably capture.^18^

Experimentally testing all naturally occurring missense variants is not feasible for most proteins due to the lack of high-throughput functional assays. To address this gap, computational tools have been developed to predict the functional impact of variants by integrating sequence conservation, structural context, physicochemical properties, and machine-learning approaches. Popular algorithms include CADD,^19^ SIFT,^20^ and PolyPhen-2,^21^, as well as newer deep-learning methods such as AlphaMissense (AM),^22^ which leverage protein structural models to improve prediction accuracy across the human proteome. Despite these advances, predictions often remain inconsistent or uncertain,^23^ particularly for proteins with complex structural features or poorly understood mechanisms, highlighting the continued need to combine *in silico* analyses with targeted experimental validation. This challenge is directly relevant to Mn transporters, where hundreds of naturally occurring variants of ZIP8, ZIP14, and ZnT10 have been documented in databases but remain largely uncharacterized. Given the essential role of these transporters in systemic Mn homeostasis and the established pathogenicity of certain mutations, systematic evaluation of their naturally occurring variants is critical to distinguish benign polymorphisms from those with potential functional or clinical significance.

In this work, we present a systematic survey of naturally occurring non-somatic missense mutations of ZIP8, ZIP14, and ZnT10, and evaluate their predicted functional consequences in the context of structural features derived from AlphaFold models. Variants with poor or inconsistent computational predictions are highlighted as priorities to guide future experimental studies aimed at clarifying the molecular mechanisms of Mn transporter dysfunction.

## Results and discussion

### Mutational landscape of Mn transporters

Before examining naturally occurring variants, we first analyzed the overall mutational landscape of human ZIP8, ZIP14, and ZnT10 using AM-predicted pathogenicity scores to obtain a global view of residue tolerance. This overview can indicate the regions where deleterious mutations are most likely to occur, which is often correlated with structural features, domain arrangements, and functional importance. To visualize these features, we mapped the pathogenicity scores onto AlphaFold-predicted structures (**Figure 1**), where red indicates residues predicted to be more sensitive, blue indicates residues predicted to be more tolerant, and white represents intermediate values. As expected, pathogenicity scores were generally correlated with secondary structure, with residues in α-helices or β-strands tending to score as more sensitive than those in flexible loops, and scores were negatively correlated with solvent or lipid exposure, reflecting the importance of buried residues for structural integrity. Remarkably, the scores aligned well with known functional features in the transport domains. For example, in ZIP8 and ZIP14, two close homologs in the ZIP family,^24, 25^ transmembrane helix 2 (TM2 or α2), TM4, TM5, and TM7 were predicted to be more sensitive than other TMs, consistent with their role in forming the metal transport pathway.^26^ In ZnT10, TM2 and TM5 showed the highest sensitivity, in agreement with their harboring of residues that constitute the high-affinity transport site.^27^ Notably, the soluble domains (extracellular domains, or ECD, for ZIP8 and ZIP4; C-terminal domain, or CTD, for ZnT10) of these transporters consistently show lower predicted pathogenicity scores than the transmembrane domains (TMDs). While this reflects the essentiality of the TMD for substrate recognition and translocation, it likely underestimates the contribution of the soluble domains to protein folding and trafficking. Indeed, our previous work has showed that several ZIP4-ECD mutations impair trafficking and cause acrodermatitis enteropathica, a rare recessive genetic disorder.^28^ Furthermore, the low pathogenicity scores in the soluble domains may be due to overlooking its role in dimerization. AM predictions are based on monomeric models, but these transporters function as dimers, as demonstrated by the structures of their homologs.^27, 29-31^

**Figure 1.**
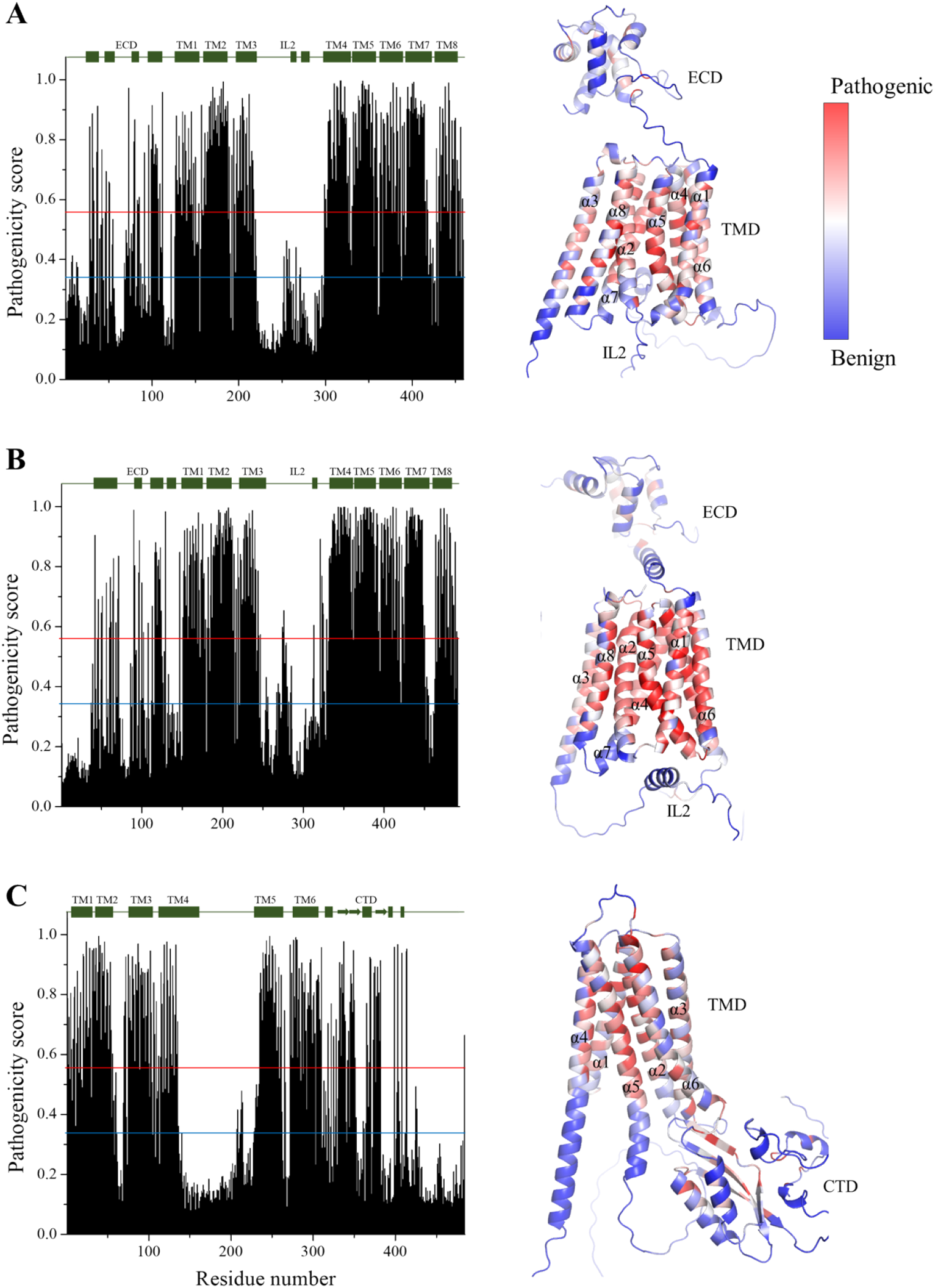
Mutational landscapes of human ZIP8 (**A**), ZIP14 (**B**), and ZnT10 (**C**). *Left*: the average pathogenicity scores predicted by AM are plotted against residue numbers. The secondary structure elements (bars for α-helices and arrows for β-strands) are indicated and labeled on top of the plots. *Right*: mapping of the average pathogenicity scores onto the AlphaFold structural models. Residues are colored according to the pathogenicity score. The red and blue lines indicate the cutoffs for pathogenic (0.564) and benign (0.34) variants, respectively.

### Variant identification and selection

To systematically catalog naturally occurring variants in ZIP8, ZIP14, and ZnT10, we used the UniProt database as our primary resource. UniProt integrates variant information retrieved from the literature and external resources, including ClinGen, ClinVar, dbSNP, gnomAD, and large-scale sequencing projects such as 1000 Genomes, ESP, ExAC, and TOPMed, providing standardized annotations of amino acid changes, allele frequencies, and functional context at the protein level. In this work, only missense variants were selected for analysis. Truncating variants (e.g., nonsense or frameshift mutations) were excluded because they are generally assumed to result in complete loss of function and thus deemed to be pathogenic. Somatic mutations (primarily retrieved from NCI-TCGA) were also excluded in order to focus the analysis on heritable variants with potential relevance to systemic Mn homeostasis. This filtering approach allowed us to compile a set of naturally occurring non-somatic missense variants suitable for functional prediction. As a result, a total of 369, 407, and 434 variants were selected for ZIP8, ZIP14, and ZnT10 (**Supplementary Material 1**), respectively.

### Pathogenicity predictions

For the selected variants, we used ProtVar, a recently developed protein-centric variant interpretation platform, to retrieve pathogenicity scores for naturally occurring missense variants of Mn transporters. ProtVar integrates CADD scores, which are composite metrics derived from integrating diverse genomic annotations to estimate variant deleteriousness,^19^ with AM predictions, which use deep learning trained on evolutionary and structural features to classify missense variants.^22^ These combined computational predictions offer different views on the potential impact of variants at the genomic and protein levels.

As shown in **Figure 2**, we compared the distributions of CADD and AM scores across the selected naturally occurring non-somatic missense variants of the three Mn transporters. Although the two prediction methods agree in ranking variant deleteriousness in general, CADD, when a loose cutoff at 20 is applied according to CADD online instruction (https://cadd.gs.washington.edu/info), consistently annotates a much larger portion of variants (56% for ZIP8, 58% for ZIP14, and 50% for ZnT10) as likely pathogenic compared to AM (**Figure 2**). This tendency of CADD to overpredict deleteriousness aligns with prior benchmarking studies showing notably higher false-positive rates: for example, more than 65% of variants in gnomAD are classified as deleterious by CADD.^32^ However, when a tight deleteriousness cutoff at 30 is applied, as suggested on the website of Ensemble (https://useast.ensembl.org/info/genome/variation/prediction/protein_function.html), CADD only annotates a very tiny fraction of variants (2.1% for ZIP8, 4.9% for ZIP14, and 5.5% for ZnT10) as pathogenic, which, as shown in the next section, fails to predict any confirmed pathogenic variants. In contrast to CADD, AM annotates much more variants as benign (73%, 68%, and 76% for ZIP8, ZIP14, and ZnT10, respectively) than pathogenic (11%, 8%, and 6%) or ambiguous (16%, 24%, and 17%). Consequently, many variants are predicted to be benign by AM but pathogenic by CADD (when the loose cutoff is applied). The most severe discrepancies can be seen in two regions in **Figure 2**: one region where CADD scores (>30) indicate very likely pathogenic variants, but AM predicts to be benign (<0.34), and another region where CADD suggests benign variants (<20), but AM marks them as pathogenic (>0.564). This discrepancy in prediction is not just limited to these two prediction tools. As shown in **Table 1**, the predictions by SIFT and PolyPhen-2 are inconsistent with those from CADD, AM, and each other without a recognizable pattern. These results indicate significant inconsistency between different computational tools, highlighting the importance of experimental validation.

**Table 1.**
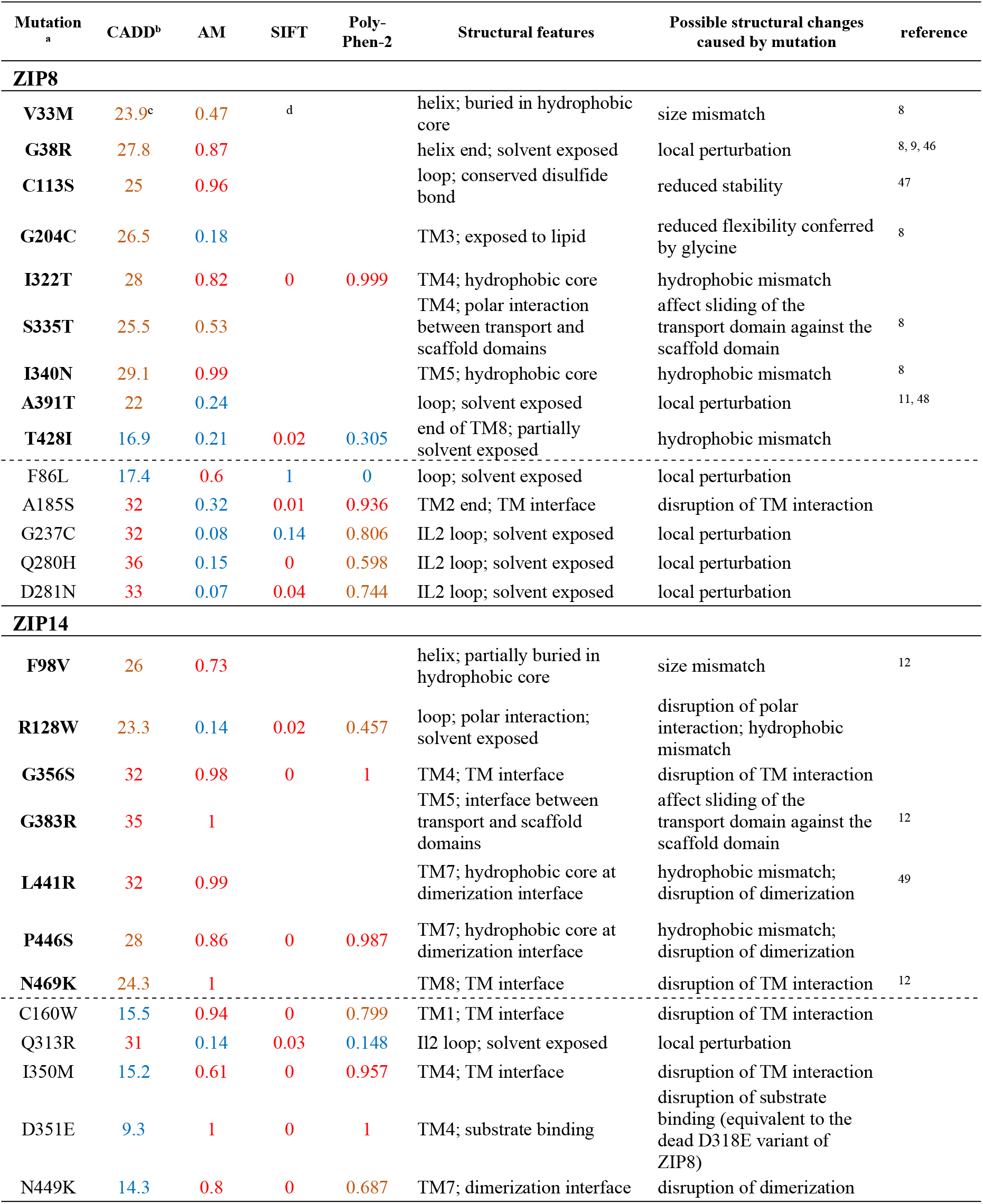
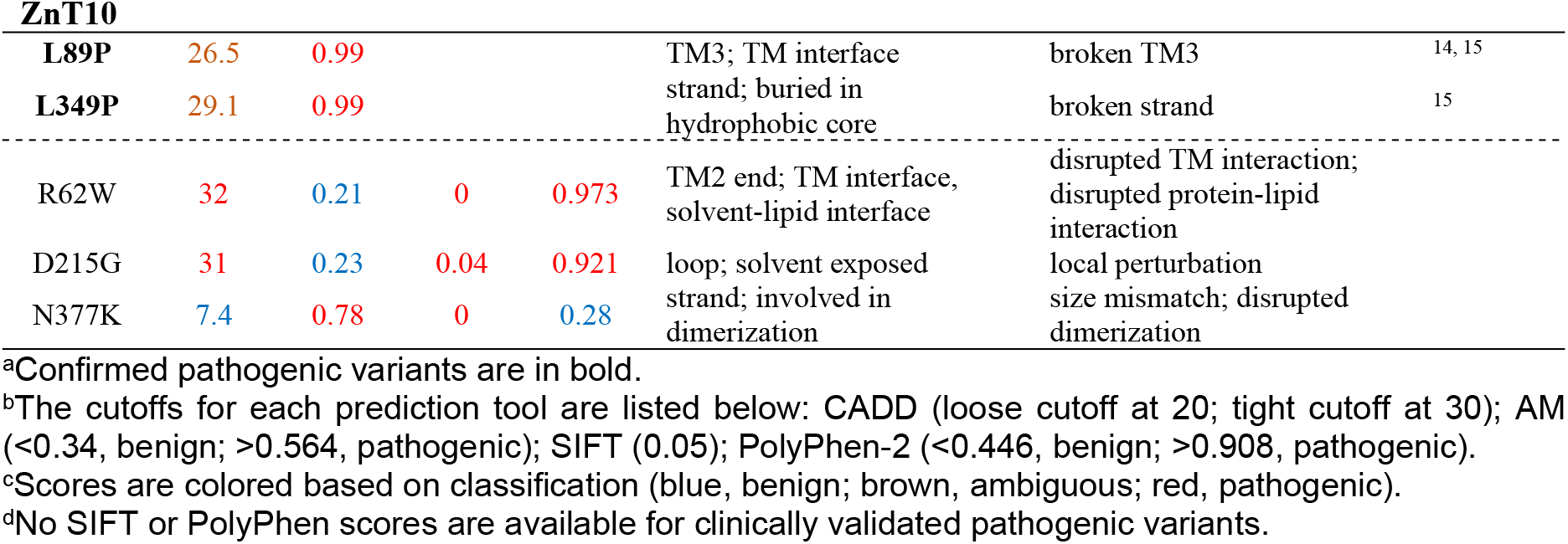
Pathogenicity predictions and structural analysis of confirmed pathogenic variants and VUS with conflicting predictions.

**Figure 2.**
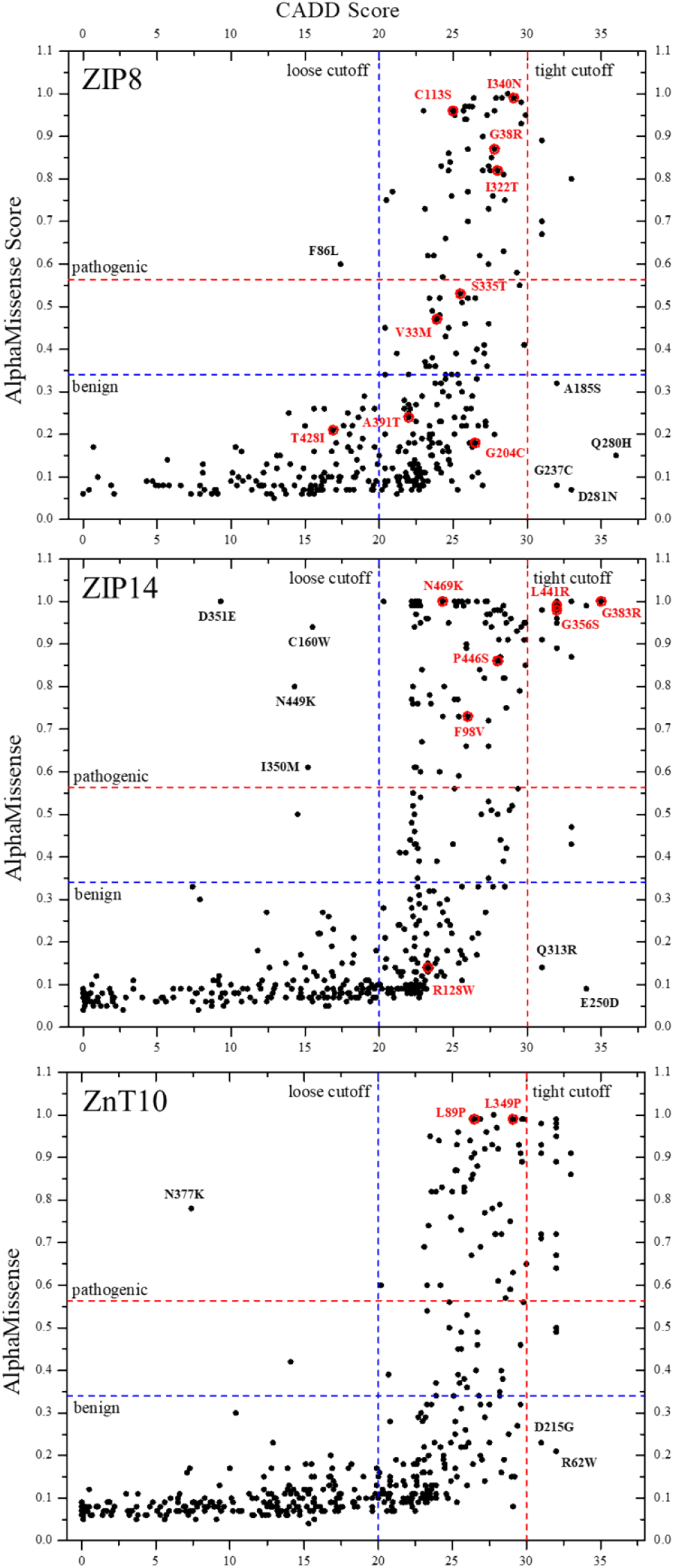
Pathogenicity predictions of naturally occurring non-somatic variants for human ZIP8, ZIP14, and ZnT10. For each selected variant, the AM score is plotted against the CADD score. The red circles indicate confirmed pathogenic variants. The dashed lines indicate the cutoffs: for CADD, the loose cutoff is set at 20, and the tight cutoff is at 30; for AM, the cutoffs for pathogenic and benign are 0.564 and 0.34, respectively. The confirmed pathogenic variants and those with drastically conflicting prediction scores are labeled.

### Comparison of pathogenicity predictions for confirmed pathogenic variants

To evaluate the predictive power of computational tools, we next compared CADD and AM scores against confirmed pathogenic annotations. In this study, a variant was considered confirmed pathogenic if either (i) there was a published report directly demonstrating that the exact mutation leads to a disease phenotype and reduced transporter function, or (ii) the variant was reported in ClinVar, which reports the relationships among human genetic variations and phenotypes,^33^ and annotated as pathogenic in UniProt, which integrates information from multiple sources. As shown in **Figure 2**, although most confirmed pathogenic variants are consistently identified by prediction tools, there remains a fraction of variants that are incorrectly annotated. This is particularly severe for ZIP8. Out of nine confirmed pathogenic variants, AM predicts only four as pathogenic, two as ambiguous, three as benign (G204C, A391T, and T428I), showing an accuracy as low as 44%. For CADD, when the tight cutoff is applied, it fails to predict any confirmed pathogenic variants; when the loose cutoff is applied, although it predicts eight as pathogenic, only missing one, and thus seemingly performs better than AM, it can be largely attributed to a large portion of variants being annotated as pathogenic (56%) under this condition. For ZIP14 and ZnT10, both CADD and AM perform better. CADD correctly predicts 100% or 38% of eight confirmed pathogenic variants of ZIP14 when a loose or tight cutoff is used, respectively, whereas AM successfully predicts seven as pathogenic, annotating one (R128W) as benign. For ZnT10, two confirmed pathogenic variants are identified by AM and CADD (when the loose cutoff is applied). Collectively, AM exhibits an accuracy of 74% for the three Mn transporters to predict pathogenicity, while the performance of CADD varies, largely depending on which cutoff is applied in analysis. These results are consistent with the study of CDKN2A, which showed that CADD achieved 45% accuracy versus ∼72% for AM,^34^ highlighting the limitations of *in silico* predictions and the need for targeted experimental validation.

### Exploration of the structural basis for prediction discrepancies

To rationalize the observed inconsistencies between CADD and AM predictions, as well as the misclassification of confirmed pathogenic variants, we examined the structural context of the relevant mutations that are labeled in **Figure 2**. These include all the confirmed pathogenic variants and the variants with drastically conflicting predictions by CADD and AM.

By mapping these variants onto AlphaFold-predicted protein structures (**Figure 3**), we obtained the structural information concerning the involved residues, aiming to identify structural features that may contribute to divergent and/or incorrect predictions. The results are summarized in **Table 1**. Our structural analysis revealed that variants occurring within α-helices or β-strands are more consistently and reliably annotated, whereas predictions are less reliable or consistent for mutations in loops or other flexible regions, including the termini of helices where conformational flexibility is higher. Notably, this pattern is consistent with the mutational landscape (**Figure 1**) where structured regions tend to be more sensitive to residue substitution than less structured regions, revealing a correlation between structural orderness and prediction accuracy for AM. Consistently, mutations affecting buried residues within the protein core tended to yield more consistent and reliable predictions than those located on solvent- or lipid-exposed surfaces. A notable example is the A391T variant in ZIP8, which is located in a solvent-exposed loop: despite its well-established pathogenicity that links it to many disease phenotypes from clinical studies,^7, 35-40^ AM predicts it as benign and CADD gives an ambiguous score. This discrepancy underscores the challenges of predicting the functional impact of variants in flexible regions, which are not uncommon to be functionally important.

**Figure 3.**
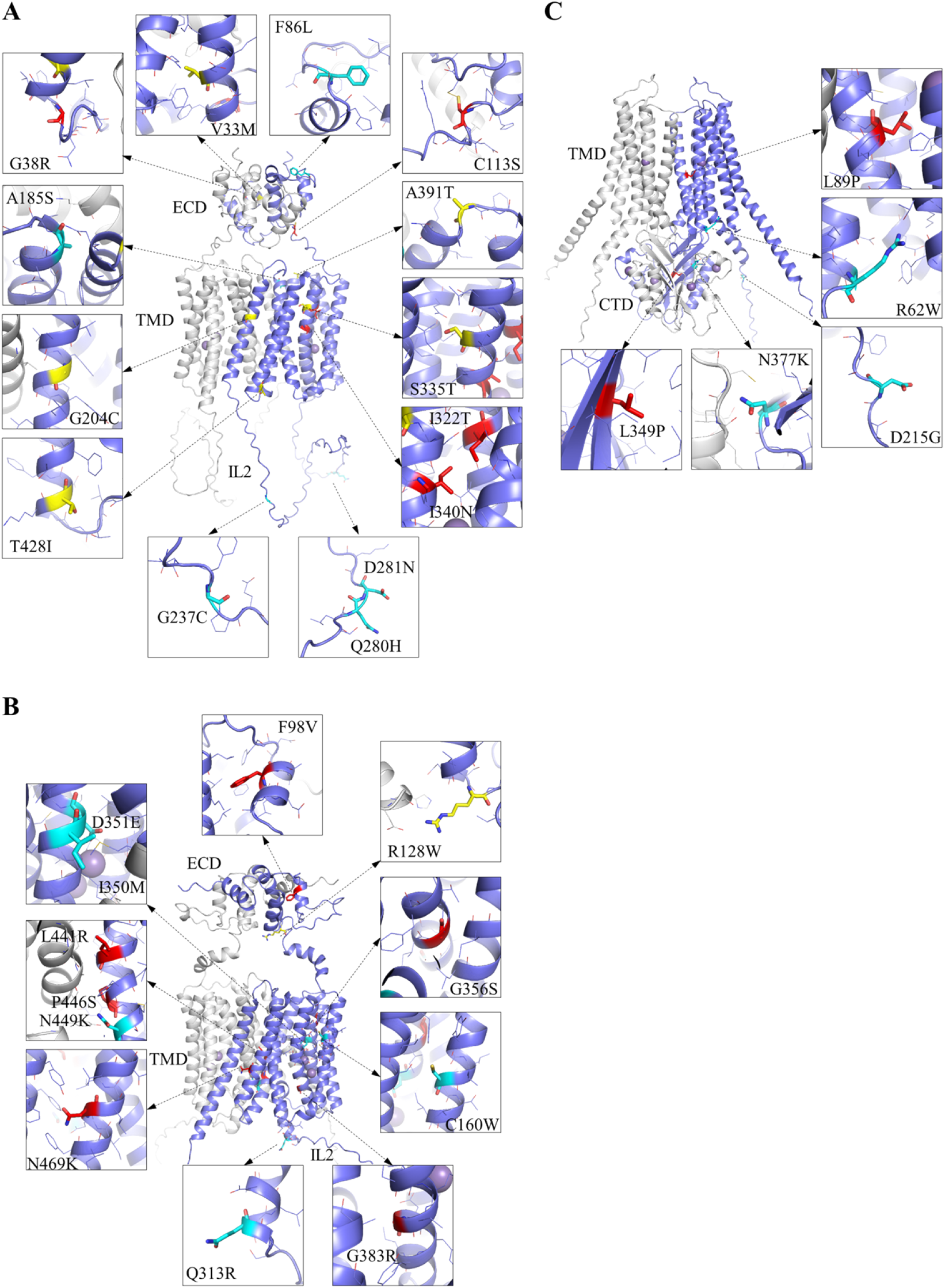
Mapping naturally occurring non-somatic missense mutations on the structural models of human ZIP8 (**A**), ZIP14 (**B**), and ZnT10 (**C**) predicted by AlphaFold 3. Only the confirmed pathogenic variants (red for correctly predicted variants by CADD and AM; yellow for variants that the computational tools failed to predict as pathogenic) and those with conflicting predictions by computational tools (cyan) are shown. Residues that are substituted are shown in stick mode and the residues within five angstroms from the changed residues are shown in line mode. The predicted dimer is shown in cartoon mode with one monomer in blue and the other in white. Manganese ions are depicted as purple spheres.

We have also proposed possible structural changes that could explain their pathogenicity. While most proposals were based primarily on local structural features, in several cases the prior knowledge regarding transport mechanism provided critical insight. For example, the S335T mutation in ZIP8, although conservative and predicted as ambiguous by both CADD and AM, is reasonably to be pathogenic because S335 lies at the interface between the transport domain and the scaffold domain. Even a small change in side-chain size, including the Ser-to-Thr substitution in this case, may disrupt the sliding of the transport domain relative to the scaffold domain, a process central to the elevator transport mechanism of ZIP proteins.^31, 41, 42^ A similar mechanistic rationale applies to the G383R mutation in ZIP14, where the introduction of a bulky, charged arginine residue at the domain interface is expected to obstruct domain movement and abolish transport activity. Likewise, the D351E substitution in ZIP14 occurs at the high-affinity metal binding site; despite being conservative, our recent work on the equivalent D318E mutation in ZIP8 demonstrated severe functional impairment,^43^ indicating that even subtle perturbations at the transport site can be deleterious. Additional examples include the L441R and P446S mutations in ZIP14, both confirmed pathogenic variants locate at the predicted dimerization interface. Since dimerization is a common structural feature of ZIP transporters,^29-31, 44, 45^ the pathogenic nature of these mutations suggests that dimerization may play a key role in transporter stability and/or function, a concept that has not yet been experimentally explored. Together, these examples highlight that structural information alone is often insufficient, and that mechanistic understanding of transporter function is indispensable for clarifying the cause of pathogenicity.

### Implications for experimental validation

Our findings underscore the importance of experimental validation, particularly because pathogenic variants such as ZIP8 A391T can be misclassified by current computational tools. If a high-throughput assay for Mn transporters is available, scanning the full spectrum of naturally occurring variants would be the most straightforward strategy, as it avoids overlooking variants that are consistently predicted to be benign by prediction tools. However, when only focused screens are feasible, priority should be given to variants with ambiguous predictions (**Supplementary Material 2)** and those with drastically conflicting results between different tools (**Figure 2**). Variants such as ZIP8 S335T also exemplify cases where mechanistic considerations strongly suggest pathogenicity despite inconclusive computational scores. Systematic follow-up of such variants through transport activity assays and protein trafficking analysis will provide critical validation.

## Conclusive remarks

In this work, we systematically analyzed naturally occurring missense variants of the three key Mn transporters, ZIP8, ZIP14, and ZnT10, using UniProt-derived variant data, computational predictions from CADD and AM, and structural context derived from AlphaFold models. Our results show that the current prediction tools frequently diverge in predictions and fail to detect even clinically validated pathogenic variants. Structural mapping and mechanistic insights proved valuable for interpreting these discrepancies, especially in cases where subtle changes at transport sites, domain interfaces, or dimerization surfaces yield strong functional consequences that are underestimated by prediction tools.

These findings emphasize that the frequently used prediction tools, while powerful, remain imperfect and must be complemented by biological knowledge and experimental validation. By identifying variants with ambiguous or conflicting predictions, we provide a prioritized set of candidates for future functional testing. Such integrative strategies are critical not only for elucidating the molecular basis of Mn transporter-related diseases but also for improving variant annotation frameworks more broadly.

## Methods

### AlphaMissense mutational landscape

The average pathogenicity scores and the corresponding structural models were retrieved from AlphaMissense website (https://alphamissense.hegelab.org/hotspot), and visualized in Pymol.

### Variant selection

All variants were retrieved from the UniProt database. Of the 506 naturally occurring variants of human ZIP8 (UniProt ID: Q9C0K1), 369 non-somatic missense variants were selected for analysis. For human ZIP14 (UniProt ID Q15043), 472 naturally occurring variants are documented; of these, 407 are non-somatic missense variants. For human ZnT10 (UniProt ID Q6XR72), 565 naturally occurring variants are documented, and 434 non-somatic missense variants were selected for analysis.

### Pathogenicity prediction

The CADD and AM scores for each selected variant were manually collected from ProVar (https://www.ebi.ac.uk/ProtVar). The SIFT and PolyPhen-2 scores were also collected from UniProt. The plots that correlating the CADD and AM scores were conducted in Origin.

### Structural model generation

AlphaFold 3 was used to generate the structural models. As the studied Mn transporters form dimers, the amino acid sequences of two polypeptide chains and multiple Mn ions were used as input. Structural models were inspected in Pymol and all structural figures were also generated in Pymol.

## Acknowledgement

This work is supported by National Institutes of Health GM140931 (to J. H.). The content is solely the responsibility of the authors and does not necessarily represent the official views of the National Institutes of Health. We thank the program “Proteins@MSU” at Michigan State University to recruit Ryan Hu to participate in this work.

## Author Contribution

R.H. and J.H. collected data, analyzed data, generated figures, and wrote manuscript.

## References

(1) Aschner, J. L.; Aschner, M. Nutritional aspects of manganese homeostasis. Molecular Aspects of Medicine 2005, 26 (4-5), 353–362. DOI: 10.1016/j.mam.2005.07.003.

(2) Chen, P.; Chakraborty, S.; Mukhopadhyay, S.; Lee, E.; Paoliello, M. M. B.; Bowman, A. B.; Aschner, M. Manganese homeostasis in the nervous system. Journal of Neurochemistry 2015, 134 (4), 601–610. DOI: 10.1111/jnc.13170.

(3) Winslow, J. W. W.; Limesand, K. H.; Zhao, N. The Functions of ZIP8, ZIP14, and ZnT10 in the Regulation of Systemic Manganese Homeostasis. Int J Mol Sci 2020, 21 (9). DOI: 10.3390/ijms21093304.

(4) Gunshin, H.; Mackenzie, B.; Berger, U. V.; Gunshin, Y.; Romero, M. F.; Boron, W. F.; Nussberger, S.; Gollan, J. L.; Hediger, M. A. Cloning and characterization of a mammalian proton-coupled metal-ion transporter. Nature 1997, 388 (6641), 482–488. DOI: 10.1038/41343.

(5) Seo, Y. A.; Li, Y.; Wessling-Resnick, M. Iron depletion increases manganese uptake and potentiates apoptosis through ER stress. NeuroToxicology 2013, 38, 67–73. DOI: 10.1016/j.neuro.2013.06.002.

(6) Mercadante, C. J.; Prajapati, M.; Conboy, H. L.; Dash, M. E.; Herrera, C.; Pettiglio, M. A.; Cintron-Rivera, L.; Salesky, M. A.; Rao, D. B.; Bartnikas, T. B. Manganese transporter Slc30a10 controls physiological manganese excretion and toxicity. Journal of Clinical Investigation 2019, 129 (12), 5442–5461. DOI: 10.1172/jci129710.

(7) Sunuwar, L.; Frkatović, A.; Sharapov, S.; Wang, Q.; Neu, H. M.; Wu, X.; Haritunians, T.; Wan, F.; Michel, S.; Wu, S.; et al. Pleiotropic ZIP8 A391T implicates abnormal manganese homeostasis in complex human disease. JCI Insight 2020, 5 (20). DOI: 10.1172/jci.insight.140978.

(8) Park, J. H.; Hogrebe, M.; Gruneberg, M.; DuChesne, I.; von der Heiden, A. L.; Reunert, J.; Schlingmann, K. P.; Boycott, K. M.; Beaulieu, C. L.; Mhanni, A. A.; et al. SLC39A8 Deficiency: A Disorder of Manganese Transport and Glycosylation. Am J Hum Genet 2015, 97 (6), 894–903. DOI: 10.1016/j.ajhg.2015.11.003.

(9) Boycott, K. M.; Beaulieu, C. L.; Kernohan, K. D.; Gebril, O. H.; Mhanni, A.; Chudley, A. E.; Redl, D.; Qin, W.; Hampson, S.; Kury, S.; et al. Autosomal-Recessive Intellectual Disability with Cerebellar Atrophy Syndrome Caused by Mutation of the Manganese and Zinc Transporter Gene SLC39A8. Am J Hum Genet 2015, 97 (6), 886–893. DOI: 10.1016/j.ajhg.2015.11.002.

(10) Bonaventura, E.; Barone, R.; Sturiale, L.; Pasquariello, R.; Alessandrì, M. G.; Pinto, A. M.; Renieri, A.; Panteghini, C.; Garavaglia, B.; Cioni, G.; et al. Clinical, molecular and glycophenotype insights in SLC39A8-CDG. Orphanet Journal of Rare Diseases 2021, 16 (1). DOI: 10.1186/s13023-021-01941-y.

(11) Lin, W.; Vann, D. R.; Doulias, P.-T.; Wang, T.; Landesberg, G.; Li, X.; Ricciotti, E.; Scalia, R.; He, M.; Hand, N. J.; et al. Hepatic metal ion transporter ZIP8 regulates manganese homeostasis and manganese-dependent enzyme activity. Journal of Clinical Investigation 2017, 127 (6), 2407–2417. DOI: 10.1172/jci90896.

(12) Tuschl, K.; Meyer, E.; Valdivia, L. E.; Zhao, N.; Dadswell, C.; Abdul-Sada, A.; Hung, C. Y.; Simpson, M. A.; Chong, W. K.; Jacques, T. S.; et al. Mutations in SLC39A14 disrupt manganese homeostasis and cause childhood-onset parkinsonism–dystonia. Nature Communications 2016, 7 (1). DOI: 10.1038/ncomms11601.

(13) Quadri, M.; Federico, A.; Zhao, T.; Breedveld Guido J.; Battisti, C.; Delnooz, C.; Severijnen, L.-A.; Di Toro Mammarella, L.; Mignarri, A.; Monti, L.; et al. Mutations in SLC30A10 Cause Parkinsonism and Dystonia with Hypermanganesemia, Polycythemia, and Chronic Liver Disease. The American Journal of Human Genetics 2012, 90 (3), 467–477. DOI: 10.1016/j.ajhg.2012.01.017.

(14) Leyva-Illades, D.; Chen, P.; Zogzas, C. E.; Hutchens, S.; Mercado, J. M.; Swaim, C. D.; Morrisett, R. A.; Bowman, A. B.; Aschner, M.; Mukhopadhyay, S. SLC30A10 Is a Cell Surface-Localized Manganese Efflux Transporter, and Parkinsonism-Causing Mutations Block Its Intracellular Trafficking and Efflux Activity. The Journal of Neuroscience 2014, 34 (42), 14079–14095. DOI: 10.1523/jneurosci.2329-14.2014.

(15) Tuschl, K.; Clayton Peter T.; Gospe Sidney M.; Gulab, S.; Ibrahim, S.; Singhi, P.; Aulakh, R.; Ribeiro Reinaldo T.; Barsottini Orlando G.; Zaki Maha S.; et al. Syndrome of Hepatic Cirrhosis, Dystonia, Polycythemia, and Hypermanganesemia Caused by Mutations in SLC30A10, a Manganese Transporter in Man. The American Journal of Human Genetics 2012, 90 (3), 457–466. DOI: 10.1016/j.ajhg.2012.01.018.

(16) Lek, M.; Karczewski, K. J.; Minikel, E. V.; Samocha, K. E.; Banks, E.; Fennell, T.; O’Donnell-Luria, A. H.; Ware, J. S.; Hill, A. J.; Cummings, B. B.; et al. Analysis of protein-coding genetic variation in 60,706 humans. Nature 2016, 536 (7616), 285–291. DOI: 10.1038/nature19057.

(17) Richards, S.; Aziz, N.; Bale, S.; Bick, D.; Das, S.; Gastier-Foster, J.; Grody, W. W.; Hegde, M.; Lyon, E.; Spector, E.; et al. Standards and guidelines for the interpretation of sequence variants: a joint consensus recommendation of the American College of Medical Genetics and Genomics and the Association for Molecular Pathology. Genetics in Medicine 2015, 17 (5), 405–424. DOI: 10.1038/gim.2015.30.

(18) Starita, L. M.; Ahituv, N.; Dunham, M. J.; Kitzman, J. O.; Roth, F. P.; Seelig, G.; Shendure, J.; Fowler, D. M. Variant Interpretation: Functional Assays to the Rescue. The American Journal of Human Genetics 2017, 101 (3), 315–325. DOI: 10.1016/j.ajhg.2017.07.014.

(19) Rentzsch, P.; Witten, D.; Cooper, G. M.; Shendure, J.; Kircher, M. CADD: predicting the deleteriousness of variants throughout the human genome. Nucleic Acids Research 2019, 47 (D1), D886–D894. DOI: 10.1093/nar/gky1016.

(20) Ng, P. C.; Henikoff, S. SIFT: predicting amino acid changes that affect protein function. Nucleic Acids Research 2003, 31 (13), 3812–3814. DOI: 10.1093/nar/gkg509.

(21) Adzhubei, I. A.; Schmidt, S.; Peshkin, L.; Ramensky, V. E.; Gerasimova, A.; Bork, P.; Kondrashov, A. S.; Sunyaev, S. R. A method and server for predicting damaging missense mutations. Nature Methods 2010, 7 (4), 248–249. DOI: 10.1038/nmeth0410-248.

(22) Cheng, J.; Novati, G.; Pan, J.; Bycroft, C.; Žemgulytė, A.; Applebaum, T.; Pritzel, A.; Wong, L. H.; Zielinski, M.; Sargeant, T.; et al. Accurate proteome-wide missense variant effect prediction with AlphaMissense. Science 2023, 381 (6664). DOI: 10.1126/science.adg7492.

(23) Kimura, H.; Lahouel, K.; Tomasetti, C.; Roberts, N. J. Functional characterization of all CDKN2A missense variants and comparison to in silico models of pathogenicity. eLife 2025, 13, RP95347. DOI: 10.7554/eLife.95347.

(24) Taylor, K. M. The LIV-1 Subfamily of Zinc Transporters: From Origins to Present Day Discoveries. Int J Mol Sci 2023, 24 (2), 1255. DOI: 10.3390/ijms24021255.

(25) Hu, J.; Jiang, Y. Evolution, classification, and mechanisms of transport, activity regulation, and substrate specificity of ZIP metal transporters. Critical Reviews in Biochemistry and Molecular Biology 2024, 59 (5), 245–266. DOI: 10.1080/10409238.2024.2405476.

(26) Zhang, T.; Liu, J.; Fellner, M.; Zhang, C.; Sui, D.; Hu, J. Crystal structures of a ZIP zinc transporter reveal a binuclear metal center in the transport pathway. Sci Adv 2017, 3 (8), e1700344. DOI: 10.1126/sciadv.1700344.

(27) Lu, M.; Fu, D. Structure of the Zinc Transporter YiiP. Science 2007, 317 (5845), 1746–1748. DOI: 10.1126/science.1143748.

(28) Kuliyev, E.; Zhang, C.; Sui, D.; Hu, J. Zinc transporter mutations linked to acrodermatitis enteropathica disrupt function and cause mistrafficking. J Biol Chem 2021, 296, 100269. DOI: 10.1016/j.jbc.2021.100269.

(29) Zhang, T.; Sui, D.; Hu, J. Structural insights of ZIP4 extracellular domain critical for optimal zinc transport. Nat Commun 2016, 7, 11979. DOI: 10.1038/ncomms11979.

(30) Pang, C.; Chai, J.; Zhu, P.; Shanklin, J.; Liu, Q. Structural mechanism of intracellular autoregulation of zinc uptake in ZIP transporters. Nat Commun 2023, 14 (1), 3404. DOI: 10.1038/s41467-023-39010-6.

(31) Zhang, Y.; Jiang, Y.; Gao, K.; Sui, D.; Yu, P.; Su, M.; Wei, G. W.; Hu, J. Structural insights into the elevator-type transport mechanism of a bacterial ZIP metal transporter. Nat Commun 2023, 14 (1), 385. DOI: 10.1038/s41467-023-36048-4.

(32) Pejaver, V.; Byrne, A. B.; Feng, B.-J.; Pagel, K. A.; Mooney, S. D.; Karchin, R.; O’Donnell-Luria, A.; Harrison, S. M.; Tavtigian, S. V.; Greenblatt, M. S.; et al. Calibration of computational tools for missense variant pathogenicity classification and ClinGen recommendations for PP3/BP4 criteria. The American Journal of Human Genetics 2022, 109 (12), 2163–2177. DOI: 10.1016/j.ajhg.2022.10.013.

(33) Landrum, M. J.; Lee, J. M.; Riley, G. R.; Jang, W.; Rubinstein, W. S.; Church, D. M.; Maglott, D. R. ClinVar: public archive of relationships among sequence variation and human phenotype. Nucleic Acids Research 2014, 42 (D1), D980–D985. DOI: 10.1093/nar/gkt1113.

(34) Kimura, H.; Lahouel, K.; Tomasetti, C.; Roberts, N. J. Functional characterization of all CDKN2A missense variants and comparison to in silico models of pathogenicity. eLife 2025, 13. DOI: 10.7554/eLife.95347.4.

(35) Ng, E.; Lind, P. M.; Lindgren, C.; Ingelsson, E.; Mahajan, A.; Morris, A.; Lind, L. Genome-wide association study of toxic metals and trace elements reveals novel associations. Human Molecular Genetics 2015, 24 (16), 4739–4745. DOI: 10.1093/hmg/ddv190.

(36) Wahlberg, K.; Arora, M.; Curtin, A.; Curtin, P.; Wright, R. O.; Smith, D. R.; Lucchini, R. G.; Broberg, K.; Austin, C. Polymorphisms in manganese transporters show developmental stage and sex specific associations with manganese concentrations in primary teeth. NeuroToxicology 2018, 64, 103–109. DOI: 10.1016/j.neuro.2017.09.003.

(37) Hill, W. D.; Marioni, R. E.; Maghzian, O.; Ritchie, S. J.; Hagenaars, S. P.; McIntosh, A. M.; Gale, C. R.; Davies, G.; Deary, I. J. A combined analysis of genetically correlated traits identifies 187 loci and a role for neurogenesis and myelination in intelligence. Molecular Psychiatry 2018, 24 (2), 169–181. DOI: 10.1038/s41380-017-0001-5.

(38) Nakata, T.; Creasey, E. A.; Kadoki, M.; Lin, H.; Selig, M. K.; Yao, J.; Lefkovith, A.; Daly, M. J.; Graham, D. B.; Xavier, R. J. A missense variant in SLC39A8 confers risk for Crohn’s disease by disrupting manganese homeostasis and intestinal barrier integrity. Proceedings of the National Academy of Sciences 2020, 117 (46), 28930–28938. DOI: 10.1073/pnas.2014742117.

(39) Verouti, S. N.; Pujol-Giménez, J.; Bermudez-Lekerika, P.; Scherler, L.; Bhardwaj, R.; Thomas, A.; Lenglet, S.; Siegrist, M.; Hofstetter, W.; Fuster, D. G.; et al. The Allelic Variant A391T of Metal Ion Transporter ZIP8 (SLC39A8) Leads to Hypotension and Enhanced Insulin Resistance. Frontiers in Physiology 2022, 13. DOI: 10.3389/fphys.2022.912277.

(40) Fujishiro, H.; Miyamoto, S.; Sumi, D.; Kambe, T.; Himeno, S. Effects of individual amino acid mutations of zinc transporter ZIP8 on manganese- and cadmium-transporting activity. Biochemical and Biophysical Research Communications 2022, 616, 26–32. DOI: 10.1016/j.bbrc.2022.05.068.

(41) Wiuf, A.; Steffen, J. H.; Becares, E. R.; Gronberg, C.; Mahato, D. R.; Rasmussen, S. G. F.; Andersson, M.; Croll, T.; Gotfryd, K.; Gourdon, P. The two-domain elevator-type mechanism of zinc-transporting ZIP proteins. Sci Adv 2022, 8 (28), eabn4331. DOI: 10.1126/sciadv.abn4331.

(42) Zhang, Y.; Jafari, M.; Zhang, T.; Sui, D.; Sagresti, L.; Merz, K. M.; Hu, J. Molecular insights into substrate translocation in an elevator-type metal transporter. Nature Communications 2024, 15 (1), 9665. DOI: 10.1038/s41467-024-54048-w.

(43) Jiang, Y.; Nikolovski, M.; Wang, T.; MacRenaris, K.; O’Halloran, T.; Hu, J. Targeting the selectivity filter to drastically alter the activity and substrate spectrum of a promiscuous metal transporter. Chemical Science 2025. DOI: 10.1039/d5sc03700j.

(44) Lin, W.; Chai, J.; Love, J.; Fu, D. Selective electrodiffusion of zinc ions in a Zrt-, Irt-like protein, ZIPB. J Biol Chem 2010, 285 (50), 39013–39020. DOI: 10.1074/jbc.M110.180620.

(45) Bin, B. H.; Fukada, T.; Hosaka, T.; Yamasaki, S.; Ohashi, W.; Hojyo, S.; Miyai, T.; Nishida, K.; Yokoyama, S.; Hirano, T. Biochemical characterization of human ZIP13 protein: a homo-dimerized zinc transporter involved in the spondylocheiro dysplastic Ehlers-Danlos syndrome. J Biol Chem 2011, 286 (46), 40255–40265. DOI: 10.1074/jbc.M111.256784.

(46) Choi, E. K.; Nguyen, T. T.; Gupta, N.; Iwase, S.; Seo, Y. A. Functional analysis of SLC39A8 mutations and their implications for manganese deficiency and mitochondrial disorders. Sci Rep 2018, 8 (1), 3163. DOI: 10.1038/s41598-018-21464-0.

(47) Riley, L. G.; Cowley, M. J.; Gayevskiy, V.; Roscioli, T.; Thorburn, D. R.; Prelog, K.; Bahlo, M.; Sue, C. M.; Balasubramaniam, S.; Christodoulou, J. A SLC39A8 variant causes manganese deficiency, and glycosylation and mitochondrial disorders. J Inherit Metab Dis 2017, 40 (2), 261–269. DOI: 10.1007/s10545-016-0010-6.

(48) Zhang, R.; Witkowska, K.; Afonso Guerra-Assunção, J.; Ren, M.; Ng, F. L.; Mauro, C.; Tucker, A. T.; Caulfield, M. J.; Ye, S. A blood pressure-associated variant of theSLC39A8gene influences cellular cadmium accumulation and toxicity. Human Molecular Genetics 2016, 25 (18), 4117–4126. DOI: 10.1093/hmg/ddw236.

(49) Bateman, J. F.; Hendrickx, G.; Borra, V. M.; Steenackers, E.; Yorgan, T. A.; Hermans, C.; Boudin, E.; Waterval, J. J.; Jansen, I. D. C.; Aydemir, T. B.; et al. Conditional mouse models support the role of SLC39A14 (ZIP14) in Hyperostosis Cranialis Interna and in bone homeostasis. PLOS Genetics 2018, 14 (4). DOI: 10.1371/journal.pgen.1007321.

